# Immunohistochemical diagnosis of *Trypanosoma vivax* in experimentally infected sheep tissues

**DOI:** 10.1101/2022.08.22.504753

**Authors:** Luiz Flávio Nepomuceno do Nascimento, Thierry Grima de Cristo, Cintia Franco, Fabiano Zanini Salbego, Renato Batista Tamanho, Renata Assis Casagrande, Joely Ferreira Figueiredo Bittar, Luiz Claudio Miletti

**Affiliations:** Laboratório de Bioquímica de Hemoparasitas e Vetores, Departamento de Produção Animal e Alimentos, Centro de Ciências Agroveterinárias, Universidade do Estado de Santa Catarina, Av. Luiz de Camões, 2090 Lages, SC, Brazil; Grupo de Patologia Veterinária, Departamento de Medicina Veterinária, Centro de Ciências Agroveterinárias, Universidade do Estado de Santa Catarina, Av. Luiz de Camões, 2090 Lages, SC, Brazil; Hospital de Clínica Veterinária, Departamento de Medicina Veterinária, Centro de Ciências Agroveterinárias, Universidade do Estado de Santa Catarina, Av. Luiz de Camões, 2090 Lages, SC, Brazil; Laboratório de Patologia Clínica e Medicina Veterinária Preventiva, Hospital Veterinário de Uberaba, Universidade de Uberaba, Av. do Tutuna, 720 Uberaba, MG, Brazil

**Keywords:** *Trypanosoma vivax*, immunohistochemistry, trypanosomiasis

## Abstract

*Trypanosoma vivax* is one of the main species responsible for animal African trypanosomiasis in West Africa and has a marked economic impact on livestock in Sub-Saharan Africa and South America endemic countries. In this work, *T. vivax* was demonstrated by immunohistochemical technique in formalin-fixed paraffin-embedded in tissues of experimentally infected sheep using polyclonal antibodies produced against formalin-fixed trypomastigotes. *T. vivax* was observed within multiple small and medium-size vessels from multiple organs, including the liver and kidneys. The immunostaining was evidenced in an intense cherry red. This is the first immunohistochemical experiment that shows *T. vivax* in fixed tissues.

## INTRODUCTION

*Trypanosoma vivax* is a flagellated protozoan belonging to the Trypanosomatidae family. It causes trypanosomosis, which may affect cattle and other ruminants’ health, leading to decreased production, growth retardation, infertility, and mortality (Dagnachew and Bezie, 2015).

The infection course of *T. vivax* depends on several factors, including the host, the presence of the vector, and the pathogenicity of the inoculum (Osório et al., 2008). In the acute phase of the disease, animals may have a fever; anemia, anorexia; apathy; progressive weight loss; lacrimation; corneal opacity, neurological and reproductive disorders (Betancur et al., 2016). In the chronic phase, clinical signs are often not observed, making diagnosis difficult and turning the animals into carriers and reservoirs of the infection (Oliveira et al., 2019).

*Trypanosoma vivax* infection can be diagnosed by parasitological, immunological, and molecular methods (Fetene et al., 2021). All these methods have advantages and disadvantages such as cross-reactions and the possibility of indicating false-negative results. The post-mortem findings can never by themselves lead to a certain diagnosis of the cause of death since there is not a single specific lesion. For an effective diagnosis, it is necessary to associate clinical data, laboratory findings, and the identification of the agent (Mandal et al., 2014).

The use of immunohistochemistry (IHC), an underexplored alternative, to identify the presence of *Trypanosoma vivax* in organs with greater tropism is considered an important *post-mortem* diagnostic test in other parasitic infection models (Reichel et al., 2007, Cooper et al., 2015, Ismail et al., 2016).

This study aimed to describe the immunohistochemical technique as a method for diagnosing *T. vivax* infection, as well as to correlate the infection with clinical and anatomopathological findings.

## MATERIALS AND METHODS

Experimental infection One male Creole sheep, eight-month-old, weighing over 40 kg were splenectomized. Their parasitological tests and PCR for *T. vivax* were negative. The animal was inoculated subcutaneously with 4 mL of a cryopreserved sample of *T. vivax* containing approximately 2×10^6^ parasites/mL. The animal was kept isolated in an enclosure fenced with mosquito nets in the sheep sector of the University of Santa Catarina State (UDESC, Lages, SC, Brazil). The animal was fed *ad libitum* with water and corn silage. Twice a day, the animal was monitored by measuring rectal temperature, globular volume, and parasitemia (fresh blood on a microscope slide/coverslip, using the WOO (Woo, 1970) and buffy coat techniques (Murray et al., 1977). This work was approved by the Animal Ethics Committee of the University of Santa Catarina State under process number 6926160818.

### Trypanosomes

At the peak of parasitemia, day 12 (2.2 × 10^5^ trypomastigotes/mL of blood), 400 mL of sheep blood were collected from the jugular vein into tubes containing EDTA. A modified protocol of that described by Cuglovici et al. (2010) was used to obtain the parasites. The blood was mixed, in an equal proportion, with a Percoll^®^ solution (Cytiva) buffered with HEPES (pH 7.4,) and centrifuged at 17,500 g at 4 ° C for 25 minutes. The parasites concentrate near the top of the gradient and were collected and stored in 15-mL centrifuge tubes.

The partially purified trypanosomes were chromatographed on a DEAE cellulose column balanced with PBS 60%/ 1.5% glucose (Lanham and Godfrey, 1970). The fractions eluted from the column were evaluated under an optical microscope and those that presented the parasites were collected and centrifuged at 6,000 *g* and 4 °C for 10 minutes. The parasites were counted in a Neubauer chamber and reached the final concentration of 1×10^8^ trypomastigotes/mL. They were kept in a 0.01 M phosphate-buffered sterile saline solution (PBS, pH 7.2), fixed with 4% formaldehyde. Subsequently, the solution was centrifuged three times at 6000 *g* and 4 °C for 5 minutes each for washing the parasites.

Production of polyclonal antisera against *Trypanosoma vivax* Antisera against *T. vivax* were produced by immunization of Wistar rats with *T. vivax* fixed with 4% formaldehyde. On day 1, rats were injected intraperitoneally with 100µL *T. vivax* fixed with 4% formaldehyde containing (1x 10^4^ parasites) mixed with 0.1 ml of Freund complete adjuvant. This injection was repeated biweekly with the same number of parasites containing Freund incomplete adjuvant instead of complete adjuvant.

The rats were anesthetized on day 52, and blood was collected by cardiac puncture. The blood of each rat was transferred into tubes without anticoagulants and centrifuged at 700 *g* for 10 minutes for serum separation. Antibody titration was assessed using the indirect immunofluorescence technique, as described by Cuglovici et al. (2010).

### Immunohistochemical method

During the necropsy, fragments of the brain, heart, lungs, kidneys, spleen, liver, skeletal muscle, abomasum, rumen, reticulum, omasum, small and large intestines, bladder, esophagus, trachea, and testicles were selected and stored in a 10% formaldehyde solution (pH 7) for a maximum of 72 hours. Sections of these organs were routinely processed for histology and embedded in paraffin for the subsequent making of 3 µm histological sections and staining using the Hematoxylin and Eosin (HE) protocol. Moreover, a histological cut of each block was made and placed on positive glass slides (*ImmunoSlide*^®^, Easypath diagnósticos, Indaiatuba, Brazil) to perform the anti-*T. vivax* Immunohistochemistry (IHC).

For histopathological evaluation, the slides were previously dewaxed in an oven at 65°C, diaphanized in xylol, rehydrated in a battery of alcohols with decreasing concentration (100% to 70%), and stained with 3% Harris’ Hematoxylin and Eosin. Afterward, staining, dehydration in a battery of alcohols with increasing concentration (80% to 100%), and diaphanization in xylol were carried out. Finally, a coverslip was mounted using a hydrophobic mounting medium (Canada balm).

The sections subjected to IHC were previously dewaxed in an oven at 65°C, diaphanized in xylol for 4 minutes, bathed twice in methanol for 2 minutes, and rehydrated in an alcohol battery with decreasing concentration (100% to 70%). For antigenic recovery, the cuts were incubated in a water bath at 100 °C for 25 minutes in a citrate buffer (pH 6). To block nonspecific reactions, skimmed milk powder was diluted in 5% phosphate-buffered saline and Tris (PBS-Tris) for 20 minutes at room temperature. The primary anti-*T. vivax* antibody was diluted 1:100 in PBS-Tris and incubated in a humid chamber at 38 °C for 1 hour and 45 minutes in a controlled temperature oven.

Subsequently, tissues were incubated with the secondary antibody, respectively an anti-rat IgG conjugated to alkaline phosphatase (1:500) (Sigma-Aldrich^®^, Merck KGaA, Darmstadt, Germany), for 60 minutes. After this, sections were developed with the Warp Red™ chromogen (Biocare Medical^®^, Pacheco, California, USA) and stained with hematoxylin for approximately 1 minute. Negative control was inserted simultaneously into the tested slides. The same section of the positive control was used. The negative was incubated with PBS instead of the primary antibody.

## RESULTS

The sera of the three rats immunized with the *Trypanosoma vivax* antigen were evaluated using the indirect immunofluorescence technique. The fluorescent parasites were visible until dilution of 1:2560 (data not shown).

*Trypanosoma vivax* was recognized readily in tissues of experimentally infected sheep stained by hematoxylin and eosin. Trypomastigote fragments, as well as entire trypomastigotes, were observed within multiple small and medium-size vessels from multiple organs, including liver, kidneys (Fig. 1. A), brain, lungs, and pancreas Trypomastigotes of *T. vivax* had an average of 18 micrometers, with pale eosinophilic cytoplasm and an elongated and predominantly basophilic central and oval nucleus with an inconspicuous nucleolus. Due to the intense cohesion between the parasites, it was not possible to observe the kinetoplast and precisely delimit the undulating membrane.

**Figure 1.**
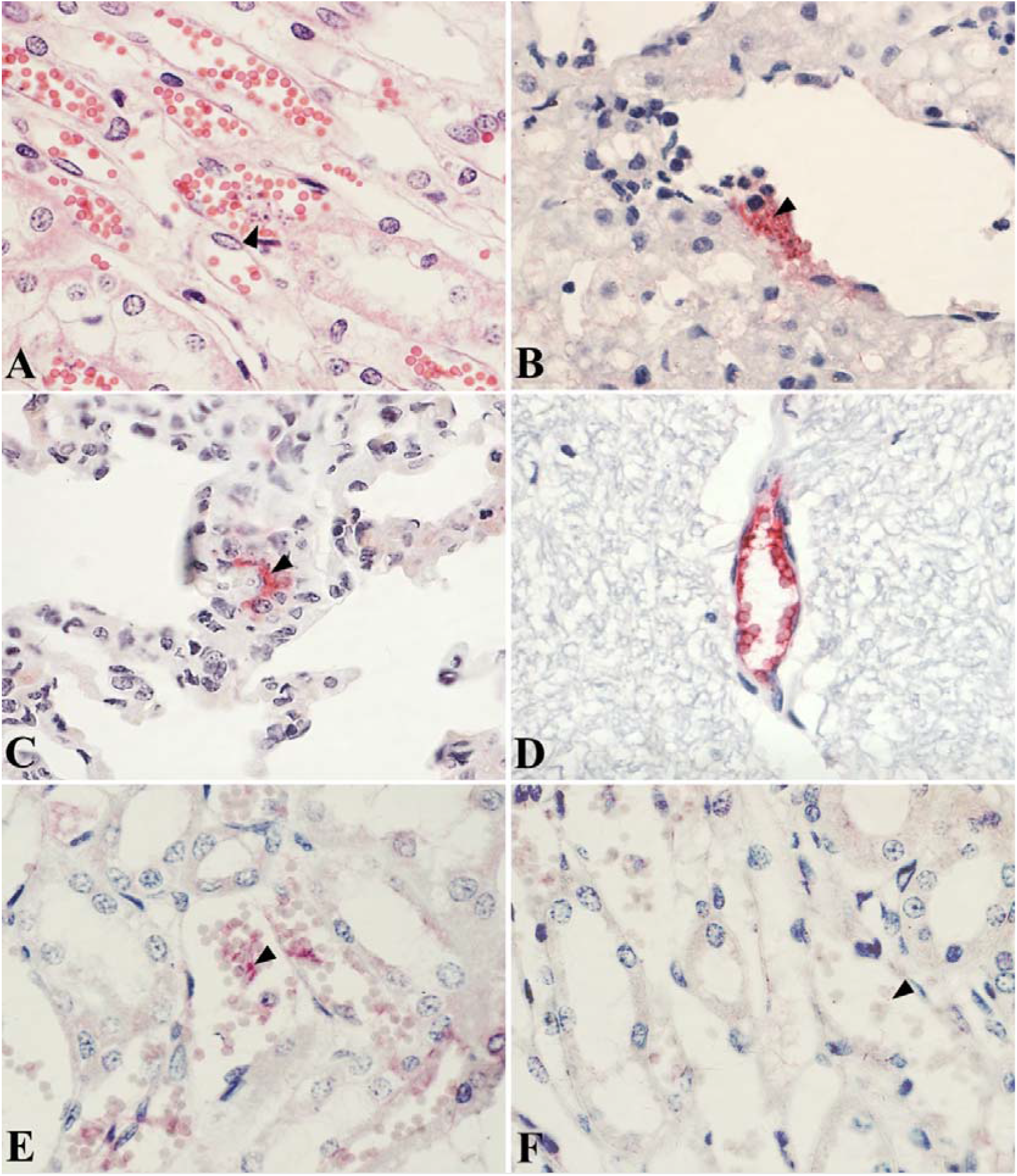
Histological and immunohistochemical evaluation of *Trypanosoma vivax* in sheep tissues. **(A)** Photomicrography of the kidneys with multiple intravascular trypomastigotes of *T. vivax* (arrowhead), stained in pale eosinophilic with an evident basophilic rounded nucleus. (Hematoxylin and eosin, ×1000). Immunohistochemical evaluation of *T. vivax* showing moderate intravascular staining in the (**B)** liver, (**C)** lung, (**D)** brain, and **(E)** kidneys (arrowheads), evidenced with intense cherry-red staining (Alkaline phosphatase bound to polymer technique, Red Warp^®^ chromogen and hematoxylin counterstain, ×1000). **(F)** Negative control made with the same kidney section as the image on the left, demonstrating trypomastigotes without immunohistochemical labeling (PBS instead of primary antibody, ×1000).

In the immunohistochemistry anti-*T. vivax*, moderate immunostaining was observed in multiple intravascular trypomastigotes, predominantly around erythrocytes and adjacent to the endothelium. This immunostaining was evidenced in an intense cherry red, and was commonly observed in the centrilobular veins of the liver (Fig. 1. B), pulmonary capillaries (Fig. 1. C), in small to medium-caliber vessels of the brain (Fig. 1. D) and with a lesser amount in the peritubular capillaries of the kidneys (Fig. 1. E). The negative control did not show any type of cherry-red immunostaining (Fig. 1. F).

## Discussion

Until recently, the pathologist has had to rely on hematoxylin and eosin or other histochemical stains and electron microscopy to show alterations that could have the involvement of infectious agents (Cartun, 1995). Immunohistochemical procedures have played an increasingly larger role in the identification of infectious disease agents in tissue sections owing to the increased availability and specificity of antibody reagents, the great sensitivity of the methods, and the relative facility with which the studies are performed. (Bacchi, et al., 1994). Furthermore, the advantages of using immunohistochemistry (IHC) for infectious disease detection include direct morphologic localization, high sensitivity that allows testing of fixed tissues and cells, and determining the presence of the parasite during chronic infection. However, it is important to note that the technique also has disadvantages. IHC stains are not standardized worldwide quantifying results is difficult and sometimes the background interferes with the analysis

The immunohistochemical technique has already been used successfully for the diagnosis of *Trypanosoma evansi* in tissues obtained from rats and water buffalos (*Bubalus bubalis*) (Sudarto et al., 1990), in tissues from naturally infected hog deer by streptavidin-biotin immunohistochemistry (Tuntasuvan et al., 2000). In an experimental infection with *Trypanosoma brucei brucei*, the immunohistochemistry revealed significant astrocytosis, loss of dendritic spines, and reduction of Purkinje cell layer of the cerebellum in Wistar rats (Adebyi et al, 2021).

The large variation in parasitemia observed in different hosts associated with non-species specific post mortem findings (FAO, 2016) demonstrates to be striking without being typical. In this work, the animal was euthanized with 12 days of infection, which is associated with an acute phase characterized by fluctuating parasitemia, fever and decreased hematocrit. As it could be demonstrated, the parasites were observed mainly in blood vessels of different tissues, which in the *post-mortem* macroscopic analysis show alterations as previously described (Uilemberg, 1998). The extravascular location of the *T. vivax* can directly damage tissues. The presence of parasites in these organs may be correlated with cases of congestion, edema, and necrosis described as clinical signs in sheep (Ogbaje et al., 2017, Batista et al., 2019) and cattle (Costa et al., 2020). It has recently been shown that *T. vivax* can also be used as a refuge for adipose tissue and skin (Machado et al., 2021). Unfortunately, these tissues were not analyzed in this work.

It is the first description of *T. vivax* in tissues of an experimental infection animal using immunohistochemistry.

